# Non-Canonical Projections from Ventral CA1 and Subicular Complex to CA3 Augments the Feedforward Hippocampal Trisynaptic Pathway

**DOI:** 10.1101/2021.02.01.429124

**Authors:** Xiaoxiao Lin, Michelle Amalraj, Crisylle Blanton, Brenda Avila, Todd C. Holmes, Douglas A. Nitz, Xiangmin Xu

## Abstract

The hippocampal formation is well documented as having a feedforward, unidirectional circuit organization termed the trisynaptic pathway. This circuit organization exists along the septotemporal axis of the hippocampal formation, but the circuit connectivity across septal to temporal regions is less well described. The emergence of viral-genetic mapping techniques enhances our ability to determine the detailed complexity of hippocampal formation circuitry. In earlier work, we mapped a subiculum back-projection to CA1 prompted by the discovery of theta wave back-propagation from the subiculum to CA1 and CA3. We reason that this circuitry may represent multiple extended non-canonical pathways involving the subicular complex and hippocampal subregions CA1 and CA3. In the present study, multiple retrograde viral tracing approaches produced robust mapping results, which supports this prediction. We find significant non-canonical synaptic inputs to dorsal hippocampal CA3 from ventral CA1, perirhinal cortex, and the subicular complex. Thus, CA1 inputs to CA3 run opposite the trisynaptic pathway and in a temporal to septal direction. Our retrograde viral tracing results are confirmed by anterograde-directed viral mapping of projections from input mapped regions to hippocampal dorsal CA3. Together, our data provide a circuit foundation to explore novel functional roles contributed by these non-canonical hippocampal circuit connections to hippocampal dynamics and behavior.

## Introduction

The extended hippocampal formation (HF) includes the entorhinal cortex (EC), the dentate gyrus (DG), hippocampus proper and the subiculum (SUB) complex. Much of the seminal work on hippocampal connectivity was carried out using conventional chemical tracing techniques [1-5]. The HF is traditionally characterized as having a feedforward, unidirectional circuit organization [6, 7]. Based on the tri-synaptic pathway model, the CA1 transfers excitatory information out of the hippocampus proper to SUB, which has been traditionally considered as a second major output stage of HF [7-13]. While this canonical HF connectivity has been well established, new viral-genetic circuit mapping approaches have revealed non-canonical HF circuits.

Our recent studies [14-16] using genetically modified rabies virus-based retrograde tracing show a significant back-projection pathway from SUB to hippocampal CA1 in the mouse. This SUB back-projection pathway (SUB-CA1) is ‘non-canonical’ as it runs directionally opposite the prominent feed-forward pathway from CA1 to SUB. Functionally, the SUB back-projection pathway contributes critically to object-place learning [14]. In contrast to the trisynaptic pathway from CA3 to CA1 to SUB, hippocampal theta frequency (8Hz) network oscillations can flow “in reverse” from SUB to CA1 and CA3 to actively modulate spike timing and local network rhythms in these subregions [17-19]. The large extent of SUB activity back-propagation suggests an even larger, non-canonical circuit network involving the subicular complex and hippocampal CA1 and CA3.

CA3 receives three major converging inputs: the mossy fiber input from DG granule cells, EC layers II/III input via the perforant path, and the extensive recurrent collateral inputs from other CA3 neurons. There is known structural heterogeneity of CA3 subregions along the transverse axis from proximal CA3 (CA3c) through middle CA3 (CA3b) to distal CA3 (CA3a). The distribution of CA3 anatomical inputs gradually changes along the transverse axis, as mossy-fiber inputs from DG taper off in the proximodistal direction. In opposite fashion, two connectivity gradients increase with associational projections between CA3 excitatory cells becoming more prominent and projections from EC layer II cells also becoming stronger along proximal to distal locations [10, 20, 21].

Gene expression markers and their distribution patterns have been used to distinguishing up to 9 CA3 domains according to their proximal/distal and septotemporal locations [22]. Domains 3, 2 and 1 correspond to the CA3a, b, and c in septal regions of CA3 as first proposed by Lorente de No. Domains 4 - 6 span approximately the mid-septotemporal half (from proximal to distal), and domains 7-9 cover the temporal pole of CA3. Physiological and functional studies reveal CA3 subregional differences corresponding to their structural and molecular expression heterogeneity along the transverse and septotemporal axes [23, 24].

While classical studies provide an overall understanding of circuit connectivity for CA3, non-canonical circuit connections have not been studied and quantitative examinations of intrinsic and extrinsic inputs (including non-canonical connections) to the subregions of CA3 are lacking. In this study, we performed multiple sets of viral tracing experiments using retrograde genetically modified canine adenovirus type 2 (CAV2), new designer variant of adenovirus associated virus (rAAV2-retro) and rabies virus, as well as anterograde herpes simplex virus (H129 strain). We find substantial non-canonical synaptic inputs to dorsal hippocampal CA3 from ventral CA1 (vCA1), perirhinal cortex (Prh), and SUB complex including ventral SUB (SUBv) and subiculum transition area (SUBtr). In particular, the input from vCA1 to dorsal CA3 runs opposite the trisynaptic pathway and opposite the direction of flow of the predominant theta frequency oscillation along the septotemporal axis. These reverse, non-canonical pathways might provide a feedback regulation for CA3 that may augment the trisynaptic circuit connections, and complement and strengthen CA3 auto-associative connections. These findings lay an anatomical foundation to consider the largely unexplored role of these non-canonical hippocampal circuit connections in hippocampal dynamics and behavior.

## Results

### Non-canonical inputs to dorsal CA3 revealed by CAV2-Cre and rAAV2-retro-Cre tracing

To map brain-wide circuit inputs to hippocampal CA3, we injected a monosynaptic retrograde viral tracer, E1-deleted canine adenovirus 2 that expresses Cre-recombinase (CAV2-ΔE1-Cre) into dorsal hippocampal CA3 (dCA3) in Cre-dependent reporter Ai9 mice for visualization of tdTomato (Fig. 1*A*). This recombinant CAV2 vector is replication incompetent due to the lack of a critical E1 gene from the viral genome and codes for retrograde expression of Cre recombinase. Based on previous studies and our own applications [14, 25, 26], this Cre expressing virus efficiently and robustly infects neurons *in vivo* and activates transgenic tdTomato expression in local and long-range connected populations of neurons in Cre-dependent reporter mouse lines.

**Figure 1.**
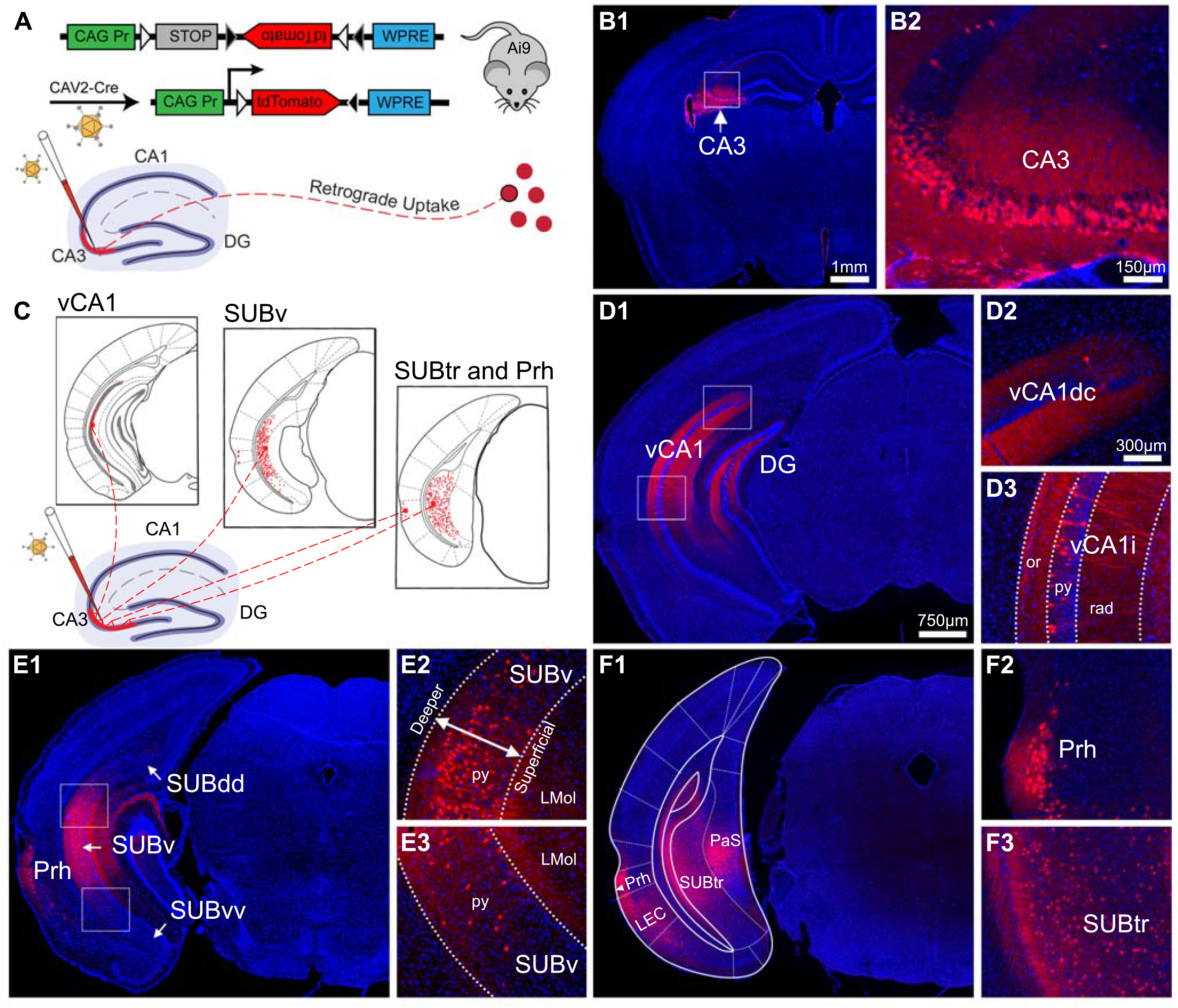
Retrograde transport of canine adenovirus 2 (CAV2-ΔE1-Cre) allows for effective mapping of canonical circuit inputs to hippocampal CA3. **A**. Strategy for retrograde monosynaptic circuit tracing using E1-deleted CAV2 viral vector expressing the Cre recombinase (CAV2-Cre) in tdTomato Cre Ai9 mice. Top: A loxP-flanked STOP cassette preventing transcription of a CAG promoter-driven red fluorescent protein (RFP) variant (tdTomato) inserted into the Gt (ROSA)26Sor locus of the Ai9 mouse genome; following Cre-mediated recombination, cells in Ai9 mice express robust tdTomato fluorescence. Bottom: CAV2-Cre injection into dorsal hippocampal CA3 leads to monosynaptic retrograde tracing of synaptic inputs to the CA3 region in the Ai9 mouse and RFP labeling near the injection site. **B**. Representative section images of viral injection sites in the hippocampal CA3. B1: The CA3 region is indicated by a white arrow. DAPI staining is blue. B2: High-magnification view of the injection site. CAV2 infected cells are labeled by tdTomato in the pyramidal cell layer of CA3. **C-H**. Results of CAV2-based retrograde monosynaptic circuit tracing from dorsal CA3 (dCA3). The labeled neurons are indicated by the white box (left, enlarged view to the right). Sources include the medial septum (MS) and vertical diagonal band (VDB) (C1 and C2), contralateral CA3 (D1 and D2), contralateral dentate hilus (E1 and E2), retromammillary nucleus (RM) (F1 and F2), median raphe nucleus (MnR) (G1 and G2) and lateral and medial entorhinal cortex (LEC and MEC) (H1 and H2). The scale bar (1 mm) applies to B1, C1, D1, E1, F1, G1 and H1; the scale bar (300 µm) applies to C2, D2, E2, F2, and H2; the scale bar (150 µm) applies to B2 and G2.

Following CAV2-Cre injection in Ai9 mice (n = 4), tdTomato expression is seen in local neurons in the dorsal CA3 injection site as well as the presynaptic neurons that provide inputs to the CA3 injection site. Note that local CAV2-Cre infected neurons are spatially restricted to the pyramidal layer of dCA3 (extending from CA3b to CA3a) without leakage to DG or CA2/CA1; thus excitatory CA3 neurons are specifically labeled (Fig. 1*B*). This is consistent with earlier findings showing that CAV2 is preferentially taken up by excitatory neurons [14, 27]. It is known that CAV2 infects epithelial cells lining the lateral ventricle (Fig. 1*B*), which we believe does not lead to unexpected results. Our results confirm that dorsal CA3 receives canonical inputs from the brain regions as identified from previous studies [3], including labeled neurons in the medial septum and vertical diagonal band (MS-DBB), contralateral CA3, dentate hilus and the entorhinal cortex (MEC and LEC) (Fig. 1*D, E* and *H*). The majority of MEC and LEC input labels are located in layer III and other deep layers. Labeled neurons are also found in the retromammillary (RM) nucleus and in the median raphe (MnR) (Fig. 1*F-G*).

Using this retrograde viral tracing approach, we additionally identify substantial non-canonical inputs to dorsal CA3, including retrogradely labeled neurons observed in vCA1 (Fig. 2*A*). We use a published gene expression marker mapping study to delineate the subregion of vCA1 along the septotemporal axis [22]. A large number of vCA1 labeled neurons are located at the pyramidal layer of intermediate vCA1 (vCA1i) (Fig. 2*B*). These pyramidal cells distribute along the deeper layer of the pyramidal cell layer and extend basal dendrites to the oriens layer (Fig. 2*B3)*. Only a few neurons are found in the upper portion of vCA1(vCA1dc) (Fig. 2*B2*).

**Figure 2:**
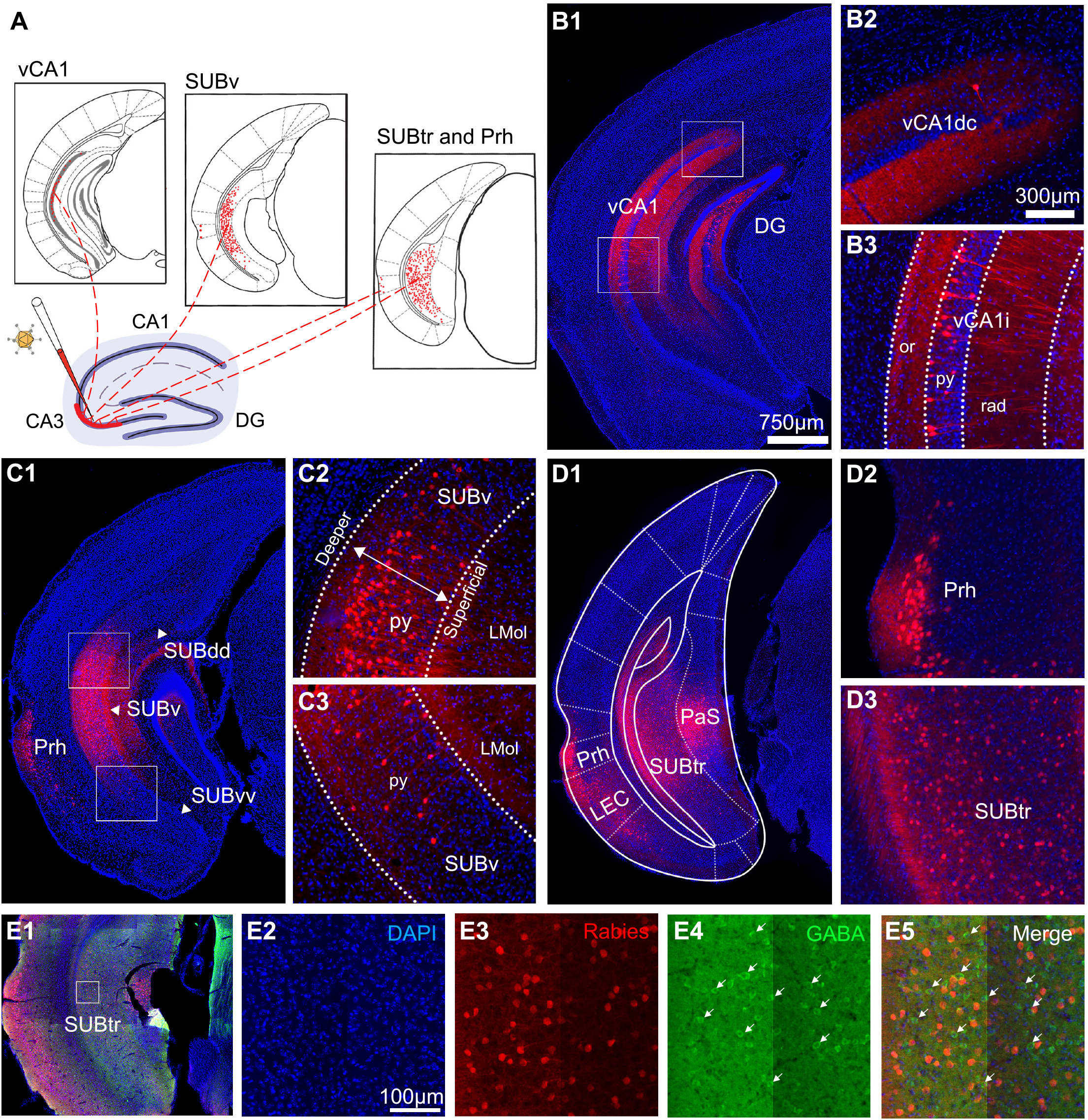
Retrograde transport of canine adenovirus 2 (CAV2-ΔE1-Cre) reveals substantial non-canonical circuit inputs from ventral hippocampal CA1 and ventral subicular complex to dorsal CA3. **A**. Results of CAV2-based retrograde monosynaptic circuit tracing from dorsal CA3, and non-canonical input mapped regions including (from left to right): ventral CA1 (vCA1), designated ventral subiculum (SUBv) and the subiculum transition area (SUBtr) and perirhinal cortex (Prh). Retrogradely labeled neurons are depicted in red. **B**. Detailed view of retrogradely labeled cells in vCA1. B1: The vCA1 input regions are indicated by white boxes. Enlarged views are shown in B2-B3. **C** and **D** are similarly formatted as in B, illustrating retrogradely labeled neurons in SUBv, SUBtr and Prh. **E**. CAV2 mapped non-GABAergic neurons in the presynaptic input regions of dCA3. E1-E5: Example images of GABA immunostaining of CAV2-labeled presynaptic neurons in SUBv. There is no colocalization between CAV2-labeled neurons and GABA immunostaining. Abbreviations: LMol, lacunosum moleculare layer of the subiculum; PaS, parasubiculum; py, pyramidal cell layer; rad, radium layer of the hippocampus; SUBdd, dorsal subiculum; SUBvv, the end tip of ventral subiculum; vCA1dc: ventral CA1, top portion; vCA1i, ventral CA1, intermediate portion.

The CAV2-Cre tracing also reveals strong circuit inputs to dCA3 from the designated ventral subiculum (SUBv) and the subiculum transition area (SUBtr) based on Franklin & Paxinos’ mouse brain atlas [28] and recent anatomical mapping studies [29-31]. As shown in the schematic Fig. 2*A*, SUBv (Bregma from -3.88mm to -4.04mm) is composed of a relatively narrow pyramidal layer and a molecular layer adjacent to the dentate gyrus. To accurately map the anatomical locations of CAV2-Cre labeled neurons, we use the SUB subregional terminology according to an earlier SUB gene expression mapping study [29] and distinguish the subicular complex subregions into the “classical” dorsal subiculum as SUBdd (aka. dSUB or SUB), the intermediate portion of the ventral subiculum as SUBv, and the end tip of the ventral subiculum as SUBvv (Fig. 2*C1*). SUBtr (Bregma from -4.16mm to -4.48mm) is close to the presubiculum (PrS), parasubiculum (PaS) and MEC.

The majority of CAV2-labeled neurons are located at the top and intermediate portions of SUBv (close to the SUBdd region). Labeled neurons are also observed at the bottom of SUBv (close to SUBvv). In comparison with the superficial pyramidal layer of SUBv, more labeled neurons are spatially localized at the deeper pyramidal layer of SUBv (Fig. 2*C*). SUBtr inputs are spatially located at the top and middle parts but are widespread with a lower density (Fig. 2*D*). The presumably excitatory input is confirmed by our immunochemical staining, as none of the CAV2-labeled cells in the SUBv and SUBtr are GABA immunopositive (Fig. 2*E*), and 90% of CAV2-labeled SUBv neurons are immuno-positive for the excitatory cell marker Ca^2+^/calmodulin-dependent protein kinase II (CaMKII). In addition to vCA1, SUBv and SUBtr inputs, we identify a direct projection from Prh to dorsal CA3 (Fig. 2*D*). Thus CAV2-Cre-mediated retrograde mapping reveals multiple non-canonical input pathways from vCA1, subregions of subicular complex (SUBv and SUBtr) and Prh to dorsal CA3 that have not been reported previously.

The finding of non-canonical inputs to dorsal CA3 by CAV2-Cre was replicated using a designer AAV variant (rAAV2-retro-Cre) that permits efficient retrograde tracing of projection neurons (Fig. 3; n = 3). Following injection into dorsal CA3 of Ai9 mice (Fig. 3*A*), both dentate granule cells and hilar cells are labeled in the rAAV2-retro-Cre experiments, while CAV2-labels only label dentate hilar cells (Fig. 3*B*). This minor variance in results indicates viral tropism differences [32-34].

**Figure 3:**
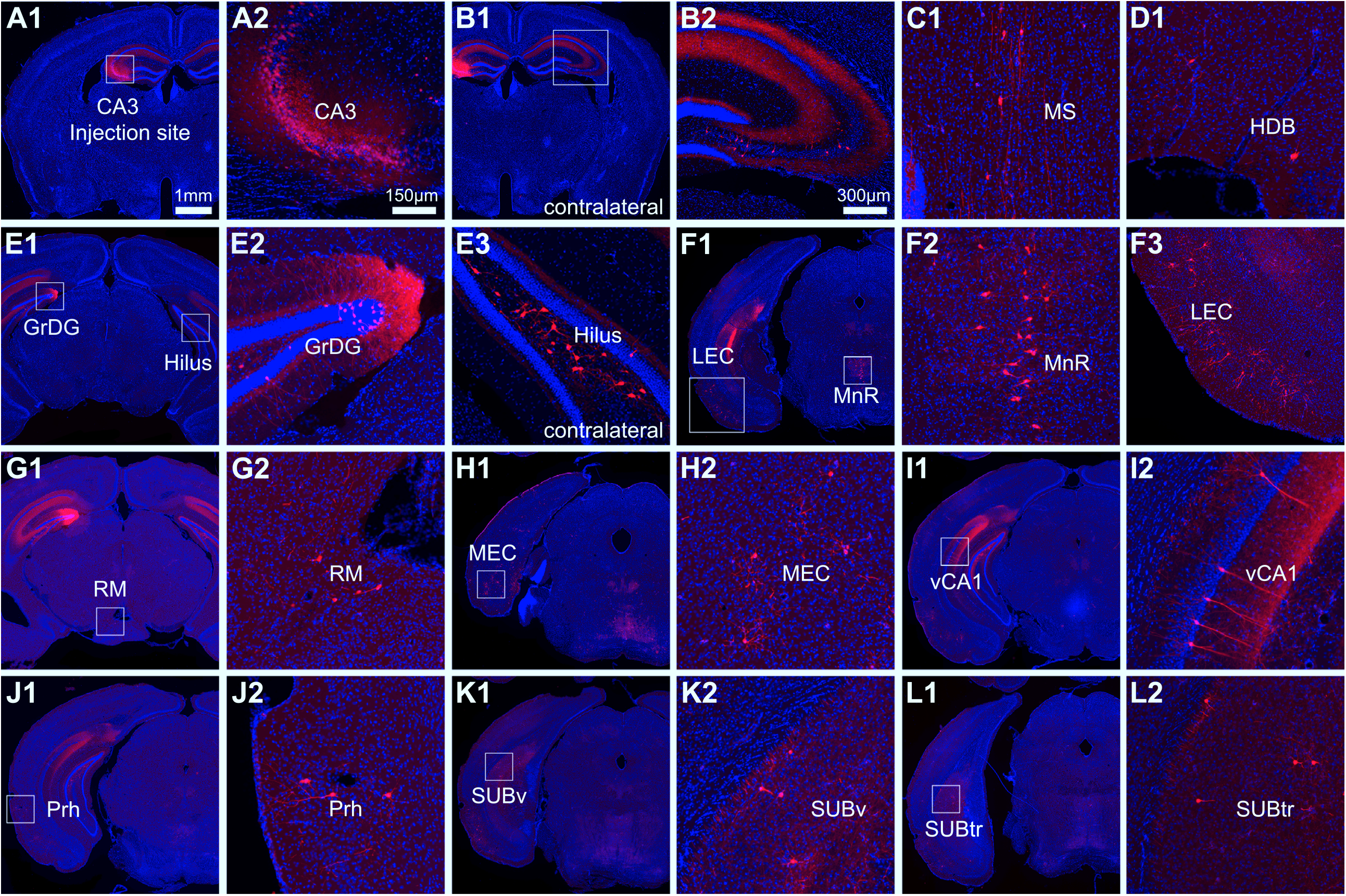
Retrograde transport of rAAV2-retro-Cre allows for effective mapping of non-canonical inputs to hippocampal CA3. **A**. Representative section images of retrograde rAAV2-retro-Cre injection site in hippocampal CA3. The CA3 injection site is indicated by a white box. DAPI staining is blue (A1). High-magnification view of the injection site (A2). **B**. The granular cell layer of the dentate gyrus (GrDG) input region and the high-magnification view of the input region. **C-F**. Sources of CA3 non-canonical afferents identified with rAAV2-retro-Cre. The labeled neurons are indicated by the white box (left, enlarged view to the right). Non-canonical input regions include the ventral CA1 (vCA1) (C1 and C2), the perirhinal cortex (Prh) (D1 and D2), the ventral subiculum (SUBv) (E1 and E2), and the subiculum transition area (SUBtr) (F1 and F2). The scale bar (1mm) in A1 applies to the low magnification images of A-F; the scale bar in A2 (150 µm) applies to high magnification images of A-F.

### Monosynaptic rabies-virus tracing of non-canonical circuit inputs to dorsal CA3

Building on CAV2-Cre and rAAV2-retro-Cre tracing experiments, we conducted monosynaptic retrograde rabies-virus-mediated tracing from dCA3. The monosynaptic rabies virus approach enables targeting of specific dCA3 subregions along the transverse axis. As rabies labeling has the feature of measurable starter cells at the injection site, connectivity strengths of input mapped brain regions can be measured quantitatively. The monosynaptic rabies viral tracing system targets specific cell types using EnvA pseudotyping and limits trans-synaptic spread to direct pre-synaptic inputs using glycoprotein gene-deleted (ΔG) rabies virus and trans complementation [15, 35, 36]. Specifically, ΔG rabies virus (deletion mutant, SAD-B19 strain) is pseudotyped with the avian sarcoma leucosis virus glycoprotein EnvA (EnvA-SADΔG rabies virus), which can only infect neurons that express avian tumor virus receptor A (TVA). TVA is an avian receptor protein that is absent in mammalian cells unless it is expressed through exogenous gene delivery. The deletion-mutant rabies virus can then be trans complemented with the expression of rabies glycoprotein (RG) in the same TVA-expressing cells to enable its retrograde spread restricted to direct presynaptic neurons. Because these presynaptic neurons lack RG expression, the virus cannot spread further beyond these cells. We used CaMKIIα-Cre (T29) mice [37] to target excitatory CA3 neurons for Cre-dependent rabies tracing. In CaMKIIα-Cre or double-transgenic mice (CaMKIIα-Cre; TVA), we virally traced circuit connections to a small population of starter CA3 pyramidal cells located in different CA3 subregions (n = 6 for CA3a, n = 5 for CA3b and n = 3 for CA3c), which were unambiguously identified by their GFP and DsRed expression from the helper AAV and ΔG-DsRed rabies genomes, respectively. Note that qualitatively similar non-canonical inputs to CA3 were shown for two slightly different approaches using CaMKIIα-Cre; TVA mice injected with the AAV8-EF1a-DIO-H2B-GFP-2A-oG and the CaMKIIα-Cre injected with the AAV8-hSyn-DIO-TC66T-2A-eGFP-2A-oG. We pooled data from these experiments for quantitative analyses of input connectivity strengths.

As shown in Fig. 4*A-D*, our rabies tracing location in more distal dCA3 is termed CA3a (corresponding to domain # 3 defined by Thompson et al. (2008) [22]). The more intermediate location of CA3 is termed CA3b (corresponding to domain # 2) and the more proximal dCA3 is termed CA3c (corresponding to domain # 1) [22]. Similar to CAV2-Cre results, the afferent circuit inputs to dCA3 subfields originate from multiple brain regions that provide canonical input, including MS-DBB, DG, ventral CA3, contralateral CA3, EC, RM and MnR (5*A-F*). Their measurements of quantitative input strengths for different dCA3 subregions are shown in Fig. 5*G-H* and Supplementary Table 1. We operationally define an input connection strength index (CSI) as the ratio of the number of presynaptic neurons in a brain region versus the number of starter neurons in the CA3. The CSI values allow us to quantitatively compare how input strengths from different brain regions to CA3 vary along the transverse axis. In addition, we calculate the proportion of inputs (PI) as the number of labeled presynaptic neurons in a brain region of interest versus the overall total labeled neurons in each case (Fig. 5 and Supplementary Table 1).

**Figure 4.**
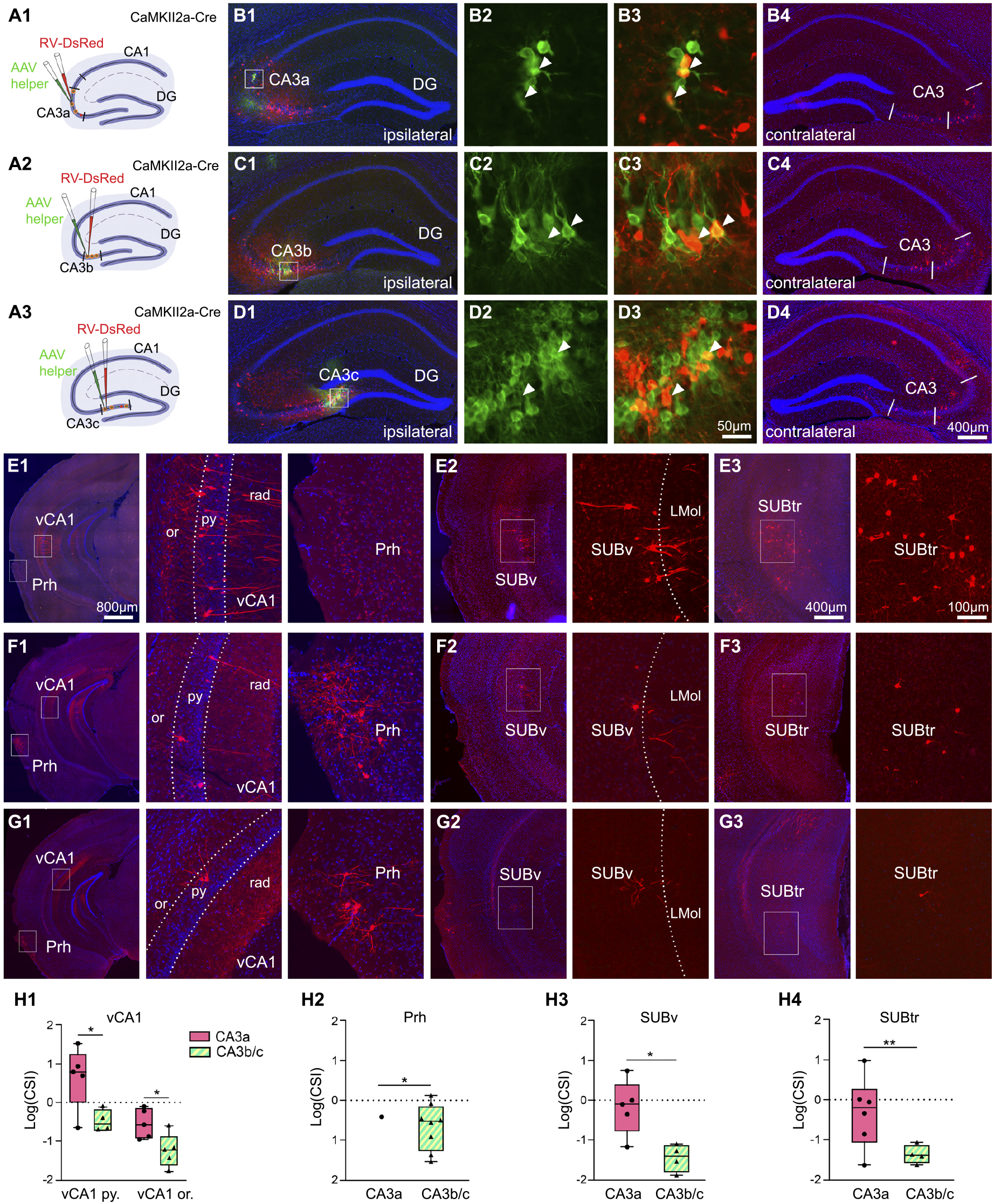
Cre-dependent retrograde monosynaptic rabies tracing of non-canonical input from vCA1, SUBv, Prh and SUBtr to CA3 varies across subregions. **A**. Schematic of retrograde monosynaptic rabies tracing from dorsal CA3 subregions in Cre transgenic mice. Injections of rabies virus (RV-DsRed: EnvA-SADΔG-RV-DsRed, red) and AAV helper virus (AAV8-hSyn-DIO-TC66T-2A-eGFP-2A-OG, green) are administered into specific subregions of dorsal CA3a-c. From top to bottom, the CA3a, CA3b, and CA3c injection sites are shown. Cre+ starter cells express both DsRed and EGFP from the rabies virus and helper AAV genomes, respectively. **B-D**. Images of injection sites of the three CA3 subregions (B1-B3 for CA3a, C1-C3 for CA3b and D1-D3 for CA3c) and corresponding rabies virus-mediated labeling of presynaptic neurons in contralateral CA3 regions with DAPI blue staining counterstain (B4, C4, and D4). The boundary of CA2 and three CA3 subregions are indicated by white lines. Injection sites are boxed in white along the transverse axis. The enlarged views of the white box regions are shown for CA3a (B3-B4), CA3b (C3-C4) and CA3c (D3-D4). Images in B3, C3 and D3 show EGFP-labeled excitatory cells (green), while the images of B4, C4 and D4 allow for visualization of DsRed and EGFP double-labeled starter neurons (white arrowheads). The images of B5, C5 and D5 show contralateral CA3 inputs for the corresponding dorsal CA3 subregions. The scale bar (400 µm) applies to B2, B5, C2, C5, D2, and D5. The scale bar (50 µm) applies to B3, B4, C3, C4, D3, and D4. **E**. Illustration of non-canonical inputs to CA3a subregions in CaMKIIα-Cre; TVA transgenic mice. E1: The boxed regions at vCA1 and Prh. These regions are shown in the middle and right panels, respectively, at a higher magnification. E2 and E3: The inputs from the SUBv and SUBtr, shown in different magnifications. **F** and **G** are formatted similarly to **E**, to illustrate non-canonical inputs to CA3b and CA3c subregions, respectively from vCA1, py, Prh, SUBv, and SUBtr. **H**. Quantitative analyses of input connection strengths measured by the log transformation of CSI across vCA1, Prh, SUBv and SUBtr following rabies tracing in CA3 subregions. vCA1 inputs are organized by the spatial location at the pyramidal layer (vCA1 py.) and oriens layer (vCA1 or.) (H1). Data from 7 CaMKIIα-Cre; TVA mice and 7 CaMKIIα-Cre mice. The data of CA3b and CA3c is combined as CA3b/c (H1-H4). Single cases are represented by black dots (circle dots for CA3a group and triangle dots for CA3b/c group). Cases with CSI = 0 are not shown on the box plot. The median of two cases is present in the Prh inputs to CA3. All other data are presented as median ± min/max; ^∗^, ^∗∗^ indicate the CSI statistical significance level of p ≤ 0.05 and p ≤ 0.01 respectively. The scale bar (800 µm) applies to E1, F1 and G1. The scale bar (400 µm) applies to E2, F2, G2, E3, F3, and G3. Scale bar (100 µm) applies to all the magnified inputs regions. Abbreviations as in Figure 1.

**Figure 5:**
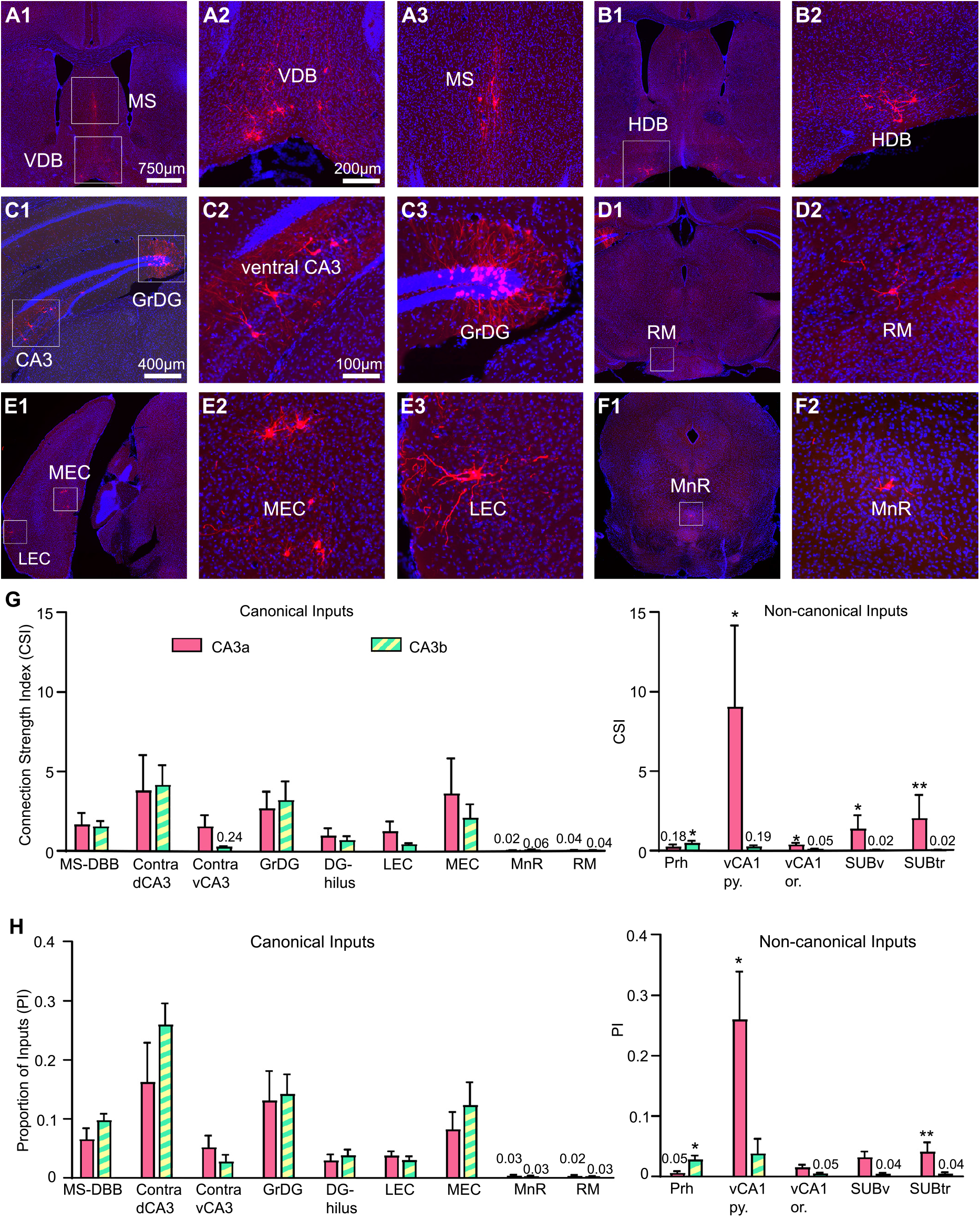
Cre-dependent monosynaptic rabies tracing of local and distant circuit connections to excitatory pyramidal neurons in subregions CA3a, CA3b, and CA3c. **A-F**. Example images showing monosynaptic rabies virus-labeled neurons from presynaptic input regions. Presynaptic cells are found in the medial septum (MS), the vertical diagonal band (VDB) (A1-A3), horizontal diagonal band (HDB) (B1-B2), ventral CA3, granule cell layer of dentate gyrus (GrDG) (C1-C3), retromammillary nucleus (RM) (D1-D2), lateral entorhinal cortex (LEC), medial entorhinal cortex (MEC) (E1-E3), and median raphe nucleus (MnR) (F1-F2). **G**. Quantitative measurements of input connection strengths measured by CSI following rabies injection in CA3a, CA3b/c. Data from 7 CaMKIIα-Cre; TVA mice and 7 CaMKIIα-Cre mice. N = 6 mice for CA3a, n = 5 for CA3b, and n = 3 for CA3c region. The data of CA3b and CA3c is combined as CA3b/c. All data are presented as mean ± SEM; ^∗^, ^∗∗^ indicate the statistical significance level of p ≤ 0.05 and p ≤ 0.01 respectively. **H**. Quantitative measurements of proportion of inputs (PI) of specific brain regions following rabies injection in CA3a, CA3b/c. The scale bar (750 µm) applies to A1, B1, D1, E1 and F1. The scale bar (200 µm) applies to A2, A3. The scale bar (400 µm) applies to C1. The scale bar (100 µm) applies to B2, C2, C3, D2, E2, E3 and F2.

To assess whether the non-canonical inputs have topographic gradients along the dCA3 transverse axis, we focus on comparing the connectivity strengths of vCA1, SUBv, SUBtr and Prh to different dCA3 subregions. We find that the more distally positioned neurons of dCA3 excitatory cells receive stronger excitatory afferent inputs from the pyramidal layer of vCA1 (vCA1 py.) compared to the more proximally positioned neurons (CSI for vCA1 py.: CA3a = 8.99 ± 5.15, n = 6; CA3b = 0.31 ± 0.13, n = 5; CA3c = 0, n = 3; CA3a vs. CA3b/c: p = 0.0386, Mann–Whitney U test) (Fig. 4*E-H* and Supplementary Table 1). The inhibitory inputs from the oriens layer of vCA1 (vCA1 or.) are also found and their CSI follows a similar trend as the pyramidal layer of vCA1 (CSI for vCA1 or.: CA3a = 0.32 ± 0.13; CA3b = 0.09 ± 0.04; CA3c = 0; CA3a vs. CA3b/c: p = 0.044, Mann–Whitney U test) (Fig. 4*E-H* and Supplementary Table 1). For direct comparison with canonical inputs, the quantitative input strengths of the non-canonical inputs for different dCA3 subregions are shown on the right side of the bar graphs (Fig. 5*G-H*).

Verifying the CAV2-Cre and rAAV2-retro-Cre tracing results, our rabies data shows that CA3 excitatory neurons receive inputs from SUBv and SUBtr regions. The superficial pyramidal neurons of the SUBv region have stronger connections with the excitatory neurons located at distal CA3 region, CA3a, compared to the ones at the proximal regions, CA3b and CA3c (CSI for SUBv: CA3a = 1.32 ± 0.87, n = 6; CA3b = 0.03 ± 0.01, n = 5; CA3c = 0, n = 3; CA3a vs. CA3b/c: p = 0.0266, Mann–Whitney U test) (Fig. 4*E-H* and Supplementary Table 1). SUBtr has a similar connectivity gradient with the subregions of CA3. CA3a received more presynaptic SUBtr inputs when compared with those received by CA3b and CA3c (CSI for SUBtr: CA3a = 1.98 ± 1.50; CA3b = 0.03 ± 0.02; CA3c = 0.01 ± 0.01; CA3a vs. CA3b/c: p = 0.008, Mann–Whitney U test) (Fig. 4*E-H* and Supplementary Table 1). In addition, we find that the Prh inputs to CA3 subregions follow an opposite gradient arrangement, in which proximal CA3, CA3c, receives the strongest Prh inputs compared to distal CA3, CA3a (CSI for Prh: CA3a = 0.18 ± 0.16, n = 6; CA3b = 0.22 ± 0.06, n = 5; CA3c = 0.75 ± 0.38, n = 3; CA3a vs. CA3b/c: p = 0.0486, Mann–Whitney U test) (Fig. 4*E-H* and Supplementary Table 1).

### Anterograde H129-mediated tracing confirms vCA1-CA3 and SUBv-CA3 projections

In order to confirm the direct projections of vCA1 and SUBv to CA3, we injected an anterograde-directed herpes virus (H129 strain; H129-G4) to vCA1 or SUBv in different cohorts of wild type C57BL/J6 mice. H129-G4 is generated by inserting binary, tandemly connected EGFP cassettes into the H129 genome. The EGFP fluorescent label of H129-G4 is sufficiently strong to visualize morphological details of labeled neurons (Fig. 6*A*). We first injected H129-G4 into the pyramidal layer of vCA1 in the C57BL/J6 mice (n = 4) and used our empirically determined 48-hour incubation time to limit the anterograde monosynaptic transmission (Fig. 6*A* and *B*). Our results show that H129-G4 labels ipsilateral and contralateral dCA3 neurons following the viral tracing from vCA1 (Fig. 6*C* and *D*). The postsynaptic neurons are located at more distal CA3 as shown in Fig. 6C2 and D2.

**Figure 6.**
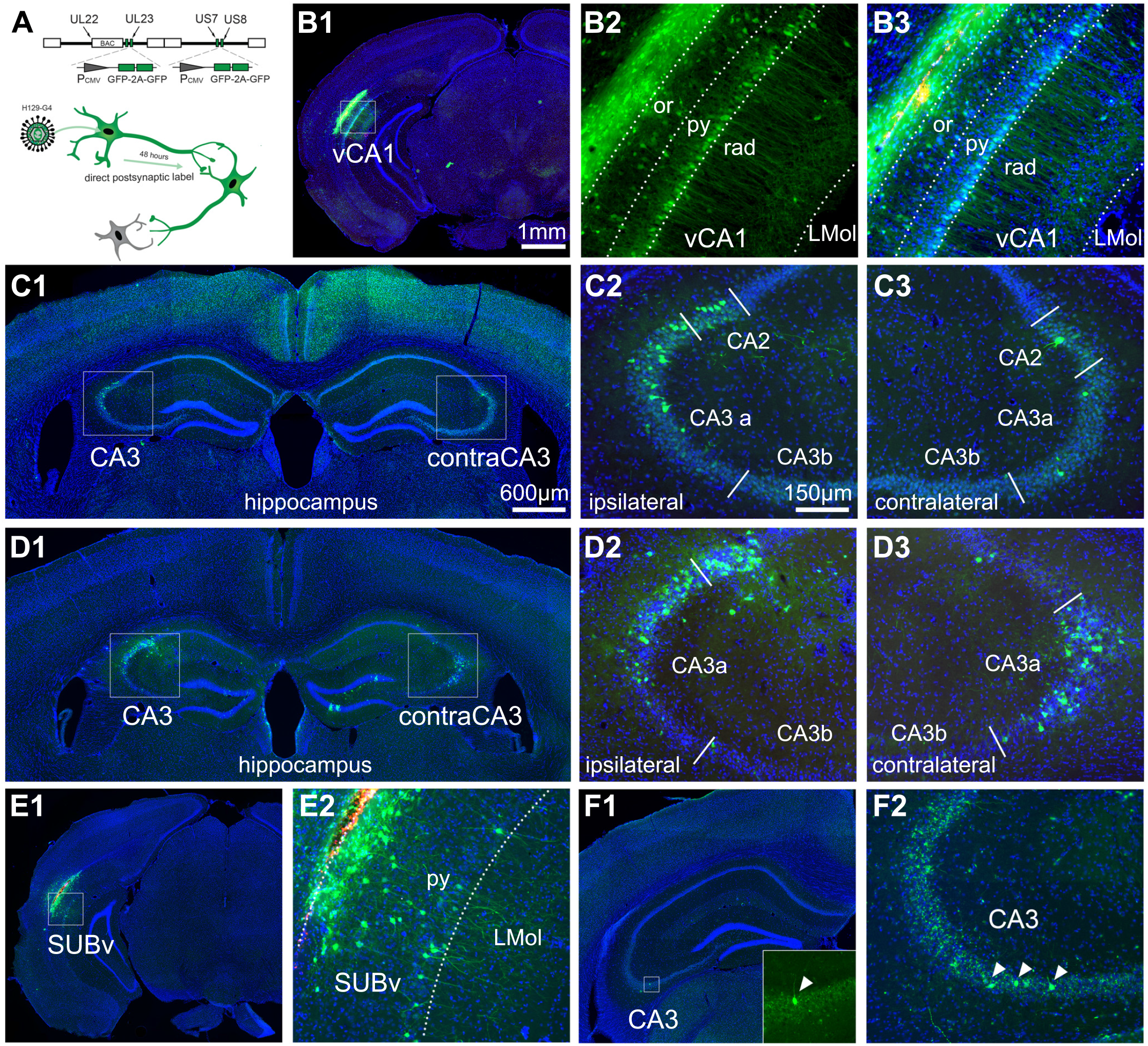
Anterograde herpes simplex virus (HSV H129) tracing verifies vCA1, SUB to dorsal CA3 projections. **A**. Schematic for anterograde H129-G4 for time-limited mapping of direct monosynaptic projections. Top: Insertion of four copies of EGFP (two tandem EGFP cassettes) into the genetically modified H129-based viral tracer. Bottom: timeline of propagation of H129-G4 after co-injection with red microspheres into the vCA1 region. **B**. Representative images of vCA1 injection site of H129-G4. B1: The H129-G4 injection site labelling (green = GFP, red = microspheres, blue = DAPI throughout all panels). B2: The EGFP-labeled neurons in the pyramidal layer (py) and oriens layer (or) of vCA1. B3: The merged image of H129-G4 expression and red microspheres that were co-injected with the H129 virus. **C**. Representative sections for H129-G4 tracing in vCA1. C1: H129-G4 labeled neurons in dorsal CA3 of both hemispheres. Enlarged views of the boxed regions in C1 are shown in C2-C3. **D**. Examples from a different case example of H129-G4 tracing from vCA1. **E-F**. Representative sections for H129-G4 tracing in SUBv. E1: The H129-G4 virus injection site. E2: shows the EGFP-labeled neurons in the py of SUBv. F1: H129-G4 labeled neurons in dorsal CA3. Enlarged view of the boxed region in F1 is shown in the bottom right corner. F2: Example from a different case of H129-G4 tracing from SUBv. The scale bar (1 mm) applies to B1 and E1; the scale bar (600 µm) applies to C1, D1 and F1; the scale bar (150 µm) applies to B2, B3, C2, C3, D2, D3, E2 and F2.

To confirm the back-projection from SUBv to CA3, a small amount of H129-G4 virus was delivered and restricted to the SUBv region (n = 3). We observe H129-G4-labeled SUBv neurons locally at the SUBv injection site (Fig. 6*E*); the number of total labeled neurons in the SUBv injection site is less than the number of labeled neurons in vCA1 (Fig. 6*B* and *E*). More sparsely labeled dorsal CA3 neurons are found following the H129-G4 injection in SUBv (Fig. 6*F*). Together with multiple anterograde and retrograde viral tracing experiments, our data demonstrate the existence of significant non-canonical circuit connections between ventral CA1 and SUB complex, and dorsal CA3, which offers the anatomical circuit basis to consider the largely unexplored functional roles of these non-canonical hippocampal circuits in hippocampal dynamics and behaviors.

## Discussion

Using multiple retrograde and anterograde viral tracers, we have identified and quantitatively mapped non-canonical circuit inputs to dorsal CA3. Unexpectedly, we discovered a prominent back-projection pathway from ventral CA1 to dorsal CA3 running opposite the trisynaptic pathway and opposite the septotemporal axis. Further, we find that non-canonical CA3 inputs include those from the subicular complex and perirhinal cortex. These non-canonical input strengths vary with dorsal CA3 locations along the transverse axis. Together our data supports the existence of an extensive, non-canonical circuitry in the HF. These non-canonical projections from vCA1 and subicular complex to CA3 augment the trisynaptic pathway through the hippocampus in a topographic fashion.

Previous studies have established the basic architecture of HF connectivity, whereas cell-type specific connections and the quantification of connectivity strengths remain less clearly revealed. We discovered non-canonical projection pathways which have not been appreciated in previous studies. Specifically, we observed projections, from vCA1, Prh, SUBv and SUBtr to dorsal CA3 excitatory neurons (Fig. 7*C*). The connectivity strength of vCA1i. SUBv, and SUBtr quantified by the ratio of presynaptic input neurons over the starter neurons in the injection site gradually decreases along the transverse axis from distal CA3 (a) to proximal CA3 (b and c). Prh projection to CA3 varies in the opposite way, increasing in strength at CA3a (Fig. 7*A-B*). Additionally, the canonical inputs from MS-DBB, DG, MnR, RM and EC regions to the pyramidal neurons of dorsal CA3 vary across subregions of CA3 along the transverse axis (Fig. 7*C*). Rabies data displayed consistent results with other retrograde viral tracers, CAV2-Cre and rAAV-retro-Cre and anterograde herpes simplex virus (H129-G4). Thus, our viral data uncovers undefined non-canonical pathways and provides alternative perspectives to understand the heterogeneity of CA3 in recent neurophysiological, genomic and functional studies.

**Figure 7.**
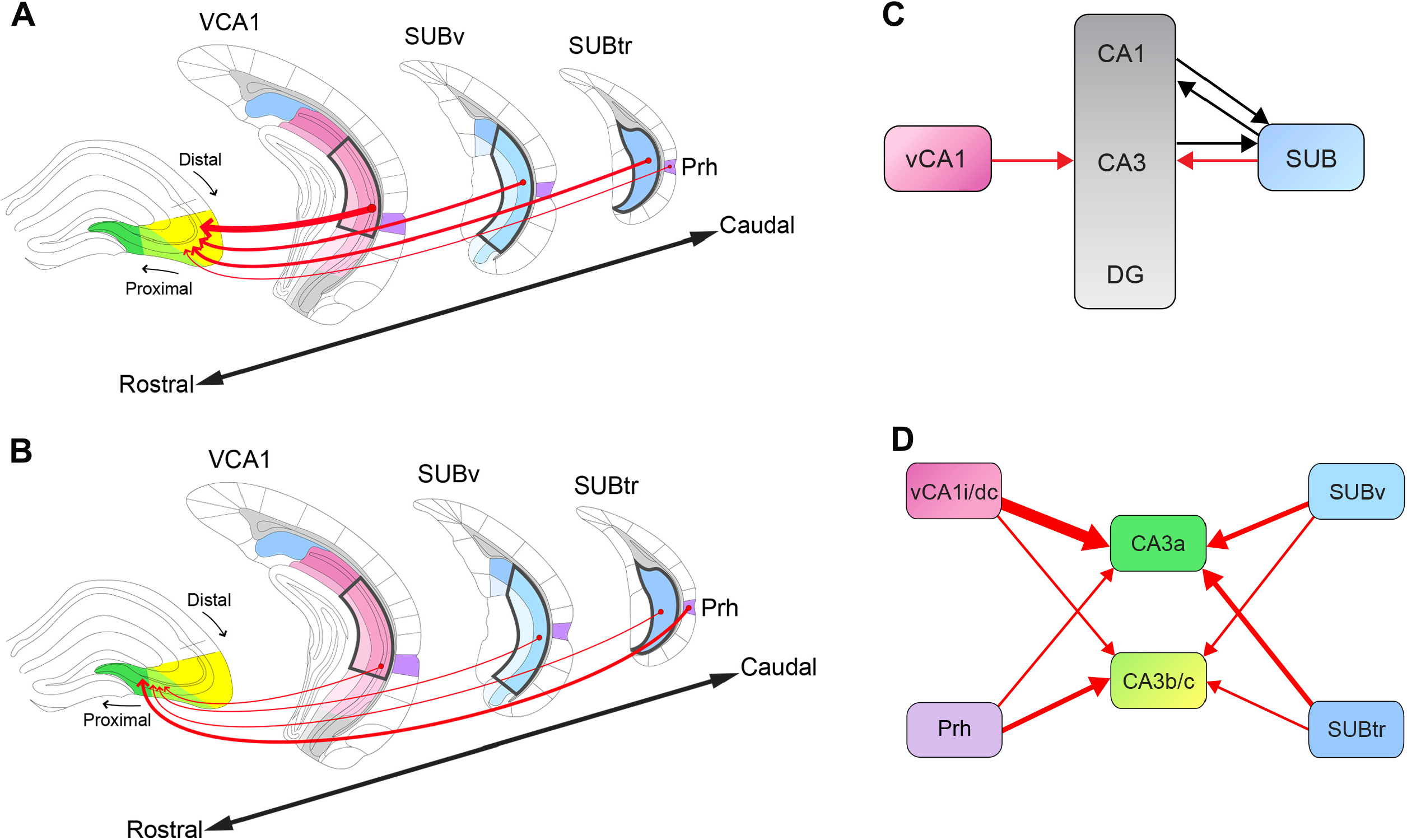
Schematic summary of canonical hippocampal circuitry and the non-canonical inputs from vCA1, SUB complex and Prh to dorsal CA3 subregions. **A**. Diagram summary of the spatial topology of non-canonical inputs to the distal region of dorsal CA3, CA3a. The input regions, vCA1, SUBv, SUBtr and Prh, are organized from the rostral to caudal direction. Gradient colors blue (the subicular complex regions) and light pink to dark pink (vCA1 subregions) represent the subregions of each input region. Shades of the same color indicate different layers in each region. Projection direction is indicated by a red arrow and the thickness of the arrow indicates the connectivity strength. **B**. The diagram is formatted similarly to A to depict the non-canonical inputs to the proximal regions of dorsal CA3, CA3b and c. **C**. The diagram depicts the novel non-canonical pathways that project in the opposite direction relative to the trisynaptic pathway. The non-canonical back-projections from SUB and vCA1 to CA3 are depicted as red lines with arrows. The previously identified feedforward projections (CA1-SUB) and non-canonical projections (SUB-CA1 (Sun et al., 2014, 18, 2019 [Ref. 14-16]), and CA3 - SUB (Ding et al., 2020 [Ref. 31]) are depicted as black lines with arrows. The vCA1 and SUB label gradients are used to indicate the subregions as matched to the corresponding colored regions in A and B. The vCA1 gradient is composed of the vCA1i and vCA1dc subregions. The SUB gradient is composed of the SUBv and SUBtr subregions. **D**. The representation diagram of differential input strengths and patterns to the dorsal CA3 subregions, CA3a and CA3b/c. Color-labeled regions are corresponding to the input regions in A and B. The vCA1i/dc gradient is composed of the colors that label the vCA1i and vCA1dc regions. The size of red arrows indicates the connectivity strength.

The differential input results of dorsal CA3 subdomains along transverse and septotemporal axes are supported by recent genomic-scale circuit mapping studies. The septal and temporal poles of the hippocampus are traditionally viewed as having a functional differentiation [38-43], but genomic approaches to defining CA3 organization also provide indication of shared functions across the septotemporal axis. The emergence of modern molecular biological techniques extends the complexity of the hippocampus according to genomic resolution and provides a molecular approach to understand circuitry and functional connections [22, 29-31, 44]. Genomic studies have divided CA3 along the transverse and septotemporal axes into 9 distinct domains, as well as CA1 and DG into 3 subdomains. In addition, recent anatomical RNA sequence studies identified 27 transcriptomic cell types residing in subiculum subdomains along the septotemporal axis [22, 29, 30]. To test our hypothesis that CA3 subregions receive distinct inputs, we used published genomic expression data to delineate CA3 subregions and related input regions including CA1 and the subiculum.

The observed tracing results show the differential projections to CA3 subdomains. Distinct gene markers divide CA1 into 3 domains CA1d, CA1i and CA1v, as dorsal, intermediate and ventral with CA1i and CA1v belong to the temporal hippocampus. Taking this perspective on organization of the hippocampus along the septotemporal axis, our data indicates CA1 efferent projections to CA3 arise from the sub-domains of CA1, CA1dc and I domains, characterized by high expression of wfs1 and Zbtb20 into distal sub-regions of CA3 (a/b).

We observed inputs to dorsal CA3 from two subregions of the subicular complex: the designated ventral subiculum (SUBv) and the subiculum transition area (SUBtr). The designated ventral subiculum (SUBv) has widespread projections to subcortical regions such as lateral septum, amygdala, bed nucleus of the striatum, hypothalamus, ventral subiculum, and lateral EC, regions implicated in reward, emotion, stress and motivation [31]. The subiculum transition area (SUBtr) topographically projects to retrosplenial cortex, parasubiculum, presubiculum, medial EC, medial mammillary nucleus and the anteroventral nucleus of thalamus, strucutures implicated in encoding of spatial location and head orientation [45-48]. Both subregions of the subicular complex project more heavily to a distal portion of dorsal CA3, overlapping with the more extensive medial EC as opposed to lateral EC inputs to dorsal CA3. This implies a role for this projection in encoding of location and orientation, but evidence for this awaits detailed electrophysiological studies of the subiculum neurons in behaving animals.

The discovery of the non-canonical pathways from vCA1i, SUBv and SUBtr extends the knowledge of hippocampal formation connectivity and its relation to learning and memory processes across the septotemporal axis. Our previous studies have shown a dorsal subiculum (SUB) back-projection to the dorsal CA1 excitatory neurons that plays an important role in facilitating object-location memory [14]. The present study reveals significant back projections from vCA1 and the subiculum to the pyramidal neurons spatially located at distal CA3. Together, these non-canonical pathways, running in the opposite direction of the traditional trisynaptic pathway, were mapped from SUB to CA1 to CA3 and to DG. These non-canonical projections therefore appear to complement the trisynaptic pathway to form a complete loop within HF.

Our data supports a new way of considering how the hippocampus processes spatial information along the septotemporal axis. The connection between the hippocampus and neocortex is topographically organized along the septotemporal axis [12]. Numerous lesion studies have shown that damage of the dorsal hippocampus impairs spatial memory, while animals with damage to the ventral hippocampus display deficits in emotional memory [39, 43, 49]. Recent physiological studies have observed theta oscillations traveling from the dorsal to ventral hippocampus [50, 51]. This implies an information processing scheme by which information from the dorsal hippocampus is integrated into the ventral regions, finally forming an output to regions such as the prefrontal cortex. In support of this, it is known that both dorsal and ventral hippocampus can represent the location of the animal in the environment, but at increasingly large scales of representation in the ventral hippocampus [52, 53]. However, the anatomical connections and functional comparison along the hippocampal septotemporal axis are still in question. Our non-canonical vCA1-dCA3, SUBv-dCA3 and SUBtr-dCA3 pathways provide evidence for direct synaptic connections between dorsal and ventral hippocampus which run opposite the direction of the trisynaptic pathway and opposite the flow of theta oscillations from septal to temporal poles. These non-canonical pathways may contribute to a crucial function relating to auto-association of CA3 via recurrent networks. In future work, it will be of interest to examine how these non-canonical inputs converging in the dorsal CA3 modulate learning and memory, and the representation of location by dorsal CA3 ensembles.

## Materials and Methods

### Animals

All experiments were conducted according to the National Institutes of Health guidelines for animal care and use, and were approved by the Institutional Animal Care and Use Committee and the Institutional Biosafety Committee of the University of California, Irvine. In the viral circuit tracing experiments, Ai9 mice expressing tdTomato fluorescence were used to study CA3 circuit connections. Transgenic CaMKIIα-Cre and CaMKIIα-Cre; TVA mice were used to map the intrinsic and extrinsic input connections of CA3 excitatory cells with genetically modified rabies virus. Ai9 mice, CaMKIIα-Cre mice and TVA mice are C57 congenic strains. Wild type C57BL/6J mice were used to verify the CA3-projecting regions with the anterograde herpes virus (H129-G4). At least 60 mice were used for the experiments, with detailed quantification performed in 27 high quality cases. See the main text for specific information.

### Viral injections

Viral injection procedure follows a previously described protocol [15]. Wild type mice were anesthetized under 1.5% isoflurane for 10 minutes with a 0.8 L/min oxygen flow rate using an isoflurane tabletop unit (HME109, Highland Medical Equipment). Mice were placed in a rodent stereotaxic frame (Leica Angle Two™ for mouse) and they were anesthetized with a continuous 1% flow of isoflurane. A small incision was made in the head, the skin reflected, and the skull exposed to show the landmarks of bregma and lambda. A three-axis micromanipulator guided by a digital atlas was used to calculate the coordinates of the injection site relative to landmarks, bregma and lambda. The virus was delivered to each subdomain of dorsal CA3 following the coordinates relative to bregma: anteroposterior (AP): -1.94mm, mediolateral (ML): -2.48mm, dorsoventral (DV): -2.24mm for CA3a; AP -1.94mm, ML: -1.97mm, DV: -2.24mm for CA3b, and AP: -2.06mm, ML: -1.92mm, DV: -2.13mm for CA3c (Fig. 4*B-D* and Supplementary Table 2). We introduced two delivery methods for viral tracers: picospritzer pressure injection and iontophoretic current injection. Pressure injection provides a large number of infected cells in the injection site, but current injection restricts the size of infected areas to spatially label the neurons in the target region. Both injection methods provide no biased inputs from other brain regions. For pressure injection, a small drill hole was made in the skull above the injection site, exposing the pia surface. A glass pipette (tip diameter, ∼20-30 μm) was loaded with the virus and then lowered into the brain at appropriate coordinates. A picospritzer (Parker Hannifin) was used to pulse viruses into the brain at a rate of 20 - 30 nl/min with 10-ms pulse duration. For iontophoresis, the virus was delivered with a positive 3-uA current at 7 s “on” and 7 s “off” cycle with a duration of 10 min. The injection pipette remained in the brain for 5 min after completion of the injection to prevent backflow of the virus. Once the injection pipette was withdrawn, the mouse was removed from the stereotaxic frame, and the incision was closed with tissue adhesive (3M Vetbond, St. Paul, MN). Mice were taken back to recover in their home cages.

#### CAV2-Cre virus

To study the circuit input connections of CA3 neurons, 0.2ul of retrograde CAV2-Cre virus (3 x 10^12^ infectious units per ml, purchased from E.J. Kremer’s group, France) was injected into the dCA3 (CA3b) of Ai9 reporter mice. After three weeks, the Ai9 mice were perfused for tissue processing.

#### rAAV2-retro-Cre virus

To confirm the CAV2-Cre results, we used retrograde rAAV2-retro-Cre virus to replicate the viral tracing experiments. 0.1ul of rAAV2-retro-hSyn-Cre virus (1.57 x 10^13^ infectious units per ml, custom packaged by Vigene Biosciences) was injected into the dCA3 (CA3b) of Ai9 reporter mice. After three weeks, the Ai9 mice were perfused for tissue processing.

#### AAV helper and rabies viruses

To map and quantitatively analyze the input strengths of CA3 pyramidal cells in the CA3 subregions, we used genetically modified monosynaptic rabies virus and transgenic mouse lines expressing Cre in CaMKIIα excitatory neurons. Rabies virus was made at the University of California, Irvine, with required cell lines and seeding viruses from E. Callaway’s group at Salk Institute for Biological Studies. For brain-wide mapping of CA3 excitatory neurons, AAV8-EF1a-DIO-H2B-GFP-2A-OG (10min, 1.54 × 10^13^ genome units per ml, custom packaged by Vigene Biosciences, Addgene plasmid #74289) was delivered through iontophoresis into target subregions in double transgenic mice of CaMKIIα-Cre; TVA. For the specific study of the intrinsic connections of CA3, AAV8-hSyn-DIO-TC66T-2A-eGFP-2A-OG (10min, 1.8 × 10^12^ genome units per ml; custom packaged by Vigene Biosciences) was delivered into the CA3 subregions of the CaMKIIα-Cre mouse line. Three weeks after the AAV injection, which allowed for the infected neurons to express high contents of RGs and GFP, the pseudotyped RG-deleted rabies virus (EnvA-SADΔG-RV-DsRed, 0.4 ul, ∼2 × 10^7^ infectious units per ml) was injected into the same target region as the AAV helper virus. The rabies virus was allowed to replicate and retrogradely spread from targeted Cre+ cell types to directly connected presynaptic cells for 9 days before the mice were perfused for tissue processing.

#### H129-G4 virus

To examine the projections of CA3 presynaptic neurons, 0.1ul of anterograde-directed herpes virus (H129-G4, 2.35 × 10^7^ genome units per ml, original reagents from the Luo lab; produced locally at the Center for Neural Circuit Mapping Center) was aimed at the following coordinates relative to bregma: anteroposterior (AP): -3.28mm, mediolateral (ML): -3.5mm, dorsoventral (DV): -3.11mm for vCA1; and AP: -4.16mm, ML: -3.22mm, DV: -3.45mm for SUBv. H129-G4 virus was allowed to replicate and anterogradely spread to postsynaptic neurons within 48 hours before the animals were perfused for tissue processing.

### Histology and immunochemical staining

The mice were perfused with 5 ml of phosphate buffered saline (PBS), followed by 25 ml PBS containing 4% paraformaldehyde. The perfused mice brains were fixed in 4% paraformaldehyde and were switched into 30% sucrose in 1 X PBS 24 hours later. In the next, the brain was frozen using dry ice and coronally sectioned in 30 µm thickness on a microtome (Leica SM2010R, Germany). One out of every three sections was mounted for examination of virally labeled neurons in different brain structures. These sections were imaged for all subsequent computer-based analyses. Some of the remaining sections were selected for neurochemical characterization of labeled cells. To identify the cell types of CAV2-labeled presynaptic neurons, GABAergic and Ca^2+^/Calmodulin-dependent protein kinase II (CaMKII) immunostaining was performed. For GABA staining, a rabbit anti-GAD65 primary antibody was used (Millipore, 1:500 dilution) followed by an Alexa Fluor 488 conjugated donkey anti-rabbit secondary antibody (Jackson ImmunoResearch, 1:200 dilution). To examine the glutamatergic labeling, selected sections were immunolabeled by a rabbit anti-CaMKII primary antibody (Sigma, 1:500 dilution) followed by the same AF488 secondary antibody used in GAD staining.

### Data Quantification

Brain slice images were acquired by using an automated slide scanning and analysis software (Metamorph, Inc.) in a high-capacity computer coupled with a fluorescent BX61 Olympus microscope and a high-sensitivity Hamamatsu CCD camera. In addition, we imaged labeled cells in selected sections with a confocal microscope (LSM 700/780, Carl Zeiss Microscopy, Nussloch, Germany) coupled with z-stack and tile scanning features under a 20X objective lens. Quantitative examinations across the series of sections were conducted for complete and unbiased analyses of virally labeled neurons by using either Metamorph or Adobe Photoshop software (Adobe Systems, San Jose, CA). For monosynaptic rabies viral tracing experiments, brain sections were examined for starter cells and presynaptic cells in different brain structures. The number of starter cells at the injection site was quantified. The mouse brain atlas was used to measure the anatomical regions of presynaptic neurons mapped by rabies virus [28]. An input connection strength index (CSI) was operationally defined as the ratio of the number of presynaptic neurons in a brain region of interest (e.g., SUB) versus the number of postsynaptic (starter) neurons in CA3. We also calculated the proportion of inputs (PI) as the ratio of the number of presynaptic inputs versus the number of total inputs. The resulting CSI and PI show similar trends for the overall input regions.

Data are presented as the mean ± SEM. For statistical comparisons between groups, the data were checked for normality of distribution. If the criteria were met, a t-test was performed to compare two groups, otherwise, a Mann-Whitney U (Wilcoxon rank-sum test) test was used. For statistical comparisons across more than two groups, one-way analysis of variance (ANOVA) or nonparametric one-way ANOVA (Kruskal–Wallis test), and related multiple comparison tests would be used for group comparisons. Alpha levels of p ≤ 0.05 were considered significant.

## Acknowledgements

This work was supported by NIH BRAIN Initiative grants [NS078434 (D.A.N, X.X.); MH120020 (X.X.)]. T.C.H. is supported by NIH R35 GM127102. The authors declare no competing interests.

---

**Supplementary Table 1.** Quantitative input strengths of CaMKIIα-Cre excitatory neurons in the CA3 subregions.

| **CA3a** | MS-DBB | Contra dCA3 | Contra vCA3 | GrDG | Hilus | LEC | MEC | MnR | RM | Prh | vCA1 py. | vCA1 or. | SUBv | SUBtr | SUBv/SUBtr |
| --- | --- | --- | --- | --- | --- | --- | --- | --- | --- | --- | --- | --- | --- | --- | --- |
| Average **CSI** | 1.6481 | 3.7765 | 1.5130 | 2.6479 | 0.9447 | 1.2193 | 3.5989 | 0.0241 | 0.0389 | 0.1815 | 8.9894 | 0.3185 | 1.3185 | 1.9762 | 2.9947 |
| SE | 0.7429 | 2.2630 | 0.7428 | 2.6479 | 0.4784 | 0.6401 | 2.2359 | 0.0182 | 0.0327 | 0.1643 | 5.1543 | 0.1281 | 0.8712 | 1.4931 | 2.4185 |
| Average **PI** | 0.0648 | 0.1620 | 0.0516 | 0.1304 | 0.0287 | 0.0370 | 0.0817 | 0.0031 | 0.0030 | 0.0053 | 0.2596 | 0.0147 | 0.0311 | 0.0405 | 0.0563 |
| SE | 0.0199 | 0.0672 | 0.0201 | 0.0511 | 0.0113 | 0.0082 | 0.0301 | 0.0020 | 0.0019 | 0.0036 | 0.0799 | 0.0052 | 0.0105 | 0.0161 | 0.0563 |
| **# of starter** | 21.8 ± 33.6 | | | | | **Total labeled neurons** | | | 1035.7 ± 1176.8 | | | | | | |

| **CA3b** | MS-DBB | Contra dCA3 | Contra vCA3 | GrDG | Hilus | LEC | MEC | MnR | RM | Prh | vCA1py. | vCA1 or. | SUBv | SUBtr | SUBv/SUBtr |
| --- | --- | --- | --- | --- | --- | --- | --- | --- | --- | --- | --- | --- | --- | --- | --- |
| Average **CSI** | 1.1124 | 4.4163 | 0.3186 | 2.0496 | 0.2994 | 0.3342 | 0.8595 | 0.0249 | 0.0357 | 0.2241 | 0.3080 | 0.0853 | 0.0321 | 0.0303 | 0.0624 |
| SE | 0.3395 | 2.0490 | 0.0819 | 1.3228 | 0.1382 | 0.1038 | 0.2720 | 0.0069 | 0.0188 | 0.0628 | 0.1292 | 0.0421 | 0.0132 | 0.0163 | 0.0271 |
| Average **PI** | 0.1055 | 0.3298 | 0.0404 | 0.1182 | 0.0310 | 0.0345 | 0.0908 | 0.0023 | 0.0024 | 0.0259 | 0.0599 | 0.0072 | 0.0062 | 0.0062 | 0.0124 |
| SE | 0.0183 | 0.0162 | 0.0171 | 0.0360 | 0.0129 | 0.0095 | 0.0286 | 0.0008 | 0.0014 | 0.0084 | 0.0377 | 0.0023 | 0.0036 | 0.0045 | 0.0080 |
| **# of starter** | 75 ± 35.5 | | | | | **Total labeled neurons** | | | 1490 ± 1374.3 | | | | | | |

| **CA3c** | MS-DBB | Contra dCA3 | Contra vCA3 | GrDG | Hilus | LEC | MEC | MnR | RM | Prh | vCA1 py. | vCA1 or. | SUBv | SUBtr | SUBv/SUBtr |
| --- | --- | --- | --- | --- | --- | --- | --- | --- | --- | --- | --- | --- | --- | --- | --- |
| Average **CSI** | 2.1953 | 3.6454 | 0.1069 | 5.0734 | 1.2361 | 0.4857 | 4.0995 | 0.1212 | 0.0541 | 0.7548 | 0.0000 | 0.0000 | 0.0000 | 0.0145 | 0.0145 |
| SE | 0.6978 | 0.9085 | 0.0180 | 2.1908 | 0.6448 | 0.2009 | 1.8492 | 0.1212 | 0.0276 | 0.3845 | 0.0000 | 0.0000 | 0.0000 | 0.0145 | 0.0145 |
| Average **PI** | 0.0828 | 0.1426 | 0.0051 | 0.1809 | 0.0480 | 0.0211 | 0.1774 | 0.0032 | 0.0023 | 0.0306 | 0.0000 | 0.0000 | 0.0000 | 0.0006 | 0.0006 |
| SE | 0.0051 | 0.0211 | 0.0020 | 0.0735 | 0.0216 | 0.0107 | 0.0948 | 0.0032 | 0.0013 | 0.0153 | 0.0000 | 0.0000 | 0.0000 | 0.0006 | 0.0006 |
| **# of starter** | 16 ± 6.2 | | | | | **Total labeled neurons** | | | 1187.7 ± 316.3 | | | | | | |

Note: The input connection strength index (CSI) is defined as the ratio of the number of presynaptic neurons in a given brain structure versus the number of starter neurons. The proportion of inputs (PI) is defined as the number of labeled neurons in a given brain structure compared to the total number of labeled neurons. The local ipsilateral CA3 input quantification is excluded. All data is presented as mean ± SEM. Abbreviations: Contra dCA3, contralateral dorsal CA3; Contra vCA3, contralateral ventral CA3; GrDG, granule cell layer of dentate gyrus; Hilus, dentate hilus; LEC, lateral entorhinal cortex; MEC, medial entorhinal cortex; MnR, median raphe nucleus; MS-DBB, medial septum, vertical diagonal band and the horizontal diagonal band; Prh, perirhinal cortex; RM, retromammillary nucleus; SUBtr, subiculum transition area; SUBv, designated ventral subiculum; vCA1 py., pyramidal cell layer of ventral CA1; vCA1 or., oriens cell layer of ventral CA1.

**Supplementary Table 2:** Mouse strains and viral injections.

| Target Region | Mouse Strain | Coordinates (mm) | Viral Injection | Method | Experiments performed |
| --- | --- | --- | --- | --- | --- |
| CA3a | CaMKIIα-Cre; TVA (N=3) | ML: -2.48; AP: -1.94; DV: -2.24 | AAV8-EF1a-DIO-H2B-GFP-2A-OG-WPRE-hGH<br>EnvA-SADΔG-RV-DsRed | Current Pressure | Retrograde monosynaptic rabies virus tracing of inputs to excitatory neurons in the dorsal CA3 subregion, CA3a |
| | CaMKIIα-Cre (N=3) | | AAV8-hSyn-DIO-TC66T-2A-eGFP-2A-OG<br>EnvA-SADΔG-RV-DsRed | Current Pressure | |
| CA3b | CaMKIIα-Cre; TVA (N=4) | ML: -1.97; AP: -1.94; DV: -2.24 | AAV8-EF1a-DIO-H2B-GFP-2A-OG-WPRE-hGH<br>EnvA-SADΔG-RV-DsRed | Current Pressure | Retrograde monosynaptic rabies virus tracing of inputs to excitatory neurons in the dorsal CA3 subregion, CA3b |
| | CaMKIIα-Cre (N=1) | | AAV8-hSyn-DIO-TC66T-2A-eGFP-2A-OG<br>EnvA-SADΔG-RV-DsRed | Current Pressure | |
| CA3c | CaMKIIα-Cre (N=3) | ML: -1.92; AP: -2.06; DV: -2.13 | AAV8-hSyn-DIO-TC66T-2A-eGFP-2A-OG<br>EnvA-SADΔG-RV-DsRed | Current Pressure | Retrograde monosynaptic rabies virus tracing of inputs to excitatory neurons in the dorsal CA3 subregion, CA3c |
| CA3 | Ai9 (RCL-tdTomato) (N=4) | ML: -1.97; AP: -1.94; DV: -2.24 | CAV2-Cre | Pressure | Retrograde CAV2-Cre tracing of excitatory inputs to the dorsal CA3. |
| CA3 | Ai9 (RCL-tdTomato) (N=3) | ML: -1.97; AP: -1.94; DV: -2.24 | rAAV2-retro-Cre | Pressure | Retrograde rAAV-retro-Cre tracing of inputs to the dorsal CA3. |
| vCA1 | C57BL/J6 (N=4) | ML: -3.5; AP: -3.28; DV: -3.11 | HSV (H129-G4) | Pressure | Anterograde HSV-mediated monosynaptic tracing verifies vCA1-CA3 projection. |
| SUBv | C57BL/J6 (N=2) | ML: -3.22; AP: -4.16; DV: -3.45 | HSV (H129-G4) | Pressure | Anterograde HSV-mediated monosynaptic tracing verifies SUBv-CA3 projection. |

Standard anterior-posterior values are relative to the bregma (see methods) are used for injections in CA3 subdomains, ventral CA1 (vCA1), and the subregion of subiculum complex (SUBv), unless otherwise specified. For all experiments, mice were either sex, ranging between 4-6 months old, and had free access to food and water in their home cages before and after surgeries.

## Reference

1. Cenquizca LA, Swanson LW. Spatial organization of direct hippocampal field CA1 axonal projections to the rest of the cerebral cortex. Brain Res Rev. 2007;56(1):1–26. doi: 10.1016/j.brainresrev.2007.05.002. PubMed PMID: 17559940; PubMed Central PMCID: PMCPMC2171036.

2. Dong HW, Swanson LW, Chen L, Fanselow MS, Toga AW. Genomic-anatomic evidence for distinct functional domains in hippocampal field CA1. Proceedings of the National Academy of Sciences of the United States of America. 2009;106(28):11794–9. doi: 10.1073/pnas.0812608106. PubMed PMID: 19561297; PubMed Central PMCID: PMCPMC2710698.

3. Witter MP. Intrinsic and extrinsic wiring of CA3: indications for connectional heterogeneity. Learning & memory. 2007;14(11):705–13. doi: 10.1101/lm.725207. PubMed PMID: 18007015.

4. Swanson LW, Sawchenko PE, Cowan WM. Evidence for collateral projections by neurons in Ammon’s horn, the dentate gyrus, and the subiculum: a multiple retrograde labeling study in the rat. The Journal of neuroscience: the official journal of the Society for Neuroscience. 1981;1(5):548–59. PubMed PMID: 6180146; PubMed Central PMCID: PMCPMC6564169.

5. Swanson LW, Cowan WM. An autoradiographic study of the organization of the efferent connections of the hippocampal formation in the rat. The Journal of comparative neurology. 1977;172(1):49–84. doi: 10.1002/cne.901720104. PubMed PMID: 65364.

6. Xu X, Sun Y, Holmes TC, Lopez AJ. Noncanonical connections between the subiculum and hippocampal CA1. The Journal of comparative neurology. 2016;524(17):3666–73. doi: 10.1002/cne.24024. PubMed PMID: 27150503; PubMed Central PMCID: PMC5050062.

7. Andersen P, Bliss TV, Lomo T, Olsen LI, Skrede KK. Lamellar organization of hippocampal excitatory pathways. Acta physiologica Scandinavica. 1969;76(1):4A–5A. doi: 10.1111/j.1748-1716.1969.tb04499.x. PubMed PMID: 5823402.

8. Scharfman HE. Evidence from simultaneous intracellular recordings in rat hippocampal slices that area CA3 pyramidal cells innervate dentate hilar mossy cells. Journal of neurophysiology. 1994;72(5):2167–80. doi: 10.1152/jn.1994.72.5.2167. PubMed PMID: 7884451.

9. MacVicar BA, Dudek FE. Local synaptic circuits in rat hippocampus: interactions between pyramidal cells. Brain research. 1980;184(1):220–3. doi: 10.1016/0006-8993(80)90602-2. PubMed PMID: 6244052.

10. Ishizuka N, Weber J, Amaral DG. Organization of intrahippocampal projections originating from CA3 pyramidal cells in the rat. The Journal of comparative neurology. 1990;295(4):580–623. doi: 10.1002/cne.902950407. PubMed PMID: 2358523.

11. de No RL. Studies on the structure of the cerebral cortex XI Continuation of the study of the ammonic system. J Psychol Neurol. 1934;46:113–77. PubMed PMID: WOS:000202445300004.

12. Steward O. Topographic organization of the projections from the entorhinal area to the hippocampal formation of the rat. The Journal of comparative neurology. 1976;167(3):285–314. doi: 10.1002/cne.901670303. PubMed PMID: 1270625.

13. Amaral DG, Kurz J. An analysis of the origins of the cholinergic and noncholinergic septal projections to the hippocampal formation of the rat. The Journal of comparative neurology. 1985;240(1):37–59. doi: 10.1002/cne.902400104. PubMed PMID: 4056104.

14. Sun Y, Jin S, Lin X, Chen L, Qiao X, Jiang L, et al. CA1-projecting subiculum neurons facilitate object-place learning. Nature neuroscience. 2019;22(11):1857–70. doi: 10.1038/s41593-019-0496-y. PubMed PMID: 31548723; PubMed Central PMCID: PMC6819262.

15. Sun Y, Nguyen AQ, Nguyen JP, L. L, Saur D, Choi J, et al. Cell-type-specific circuit connectivity of hippocampal CA1 revealed through Cre-dependent rabies tracing. Cell reports. 2014;7(1):269–80. doi: 10.1016/j.celrep.2014.02.030. PubMed PMID: 24656815; PubMed Central PMCID: PMC3998524.

16. Sun Y, Nitz DA, Holmes TC, Xu X. Opposing and Complementary Topographic Connectivity Gradients Revealed by Quantitative Analysis of Canonical and Noncanonical Hippocampal CA1 Inputs. eNeuro. 2018;5(1). doi: 10.1523/ENEURO.0322-17.2018. PubMed PMID: 29387780; PubMed Central PMCID: PMC5790753.

17. Harris E, Stewart M. Propagation of synchronous epileptiform events from subiculum backward into area CA1 of rat brain slices. Brain research. 2001;895(1-2):41–9. doi: 10.1016/s0006-8993(01)02023-6. PubMed PMID: 11259758.

18. Jackson J, Amilhon B, Goutagny R, Bott JB, Manseau F, Kortleven C, et al. Reversal of theta rhythm flow through intact hippocampal circuits. Nature neuroscience. 2014;17(10):1362–70. doi: 10.1038/nn.3803. PubMed PMID: 25174002.

19. Wozny C, Knopp A, Lehmann TN, Heinemann U, Behr J. The subiculum: a potential site of ictogenesis in human temporal lobe epilepsy. Epilepsia. 2005;46 Suppl 5:17–21. doi: 10.1111/j.1528-1167.2005.01066.x. PubMed PMID: 15987248.

20. Ishizuka N, Cowan WM, Amaral DG. A quantitative analysis of the dendritic organization of pyramidal cells in the rat hippocampus. The Journal of comparative neurology. 1995;362(1):17–45. doi: 10.1002/cne.903620103. PubMed PMID: 8576427.

21. Li XG, Somogyi P, Ylinen A, Buzsaki G. The hippocampal CA3 network: an in vivo intracellular labeling study. The Journal of comparative neurology. 1994;339(2):181–208. doi: 10.1002/cne.903390204. PubMed PMID: 8300905.

22. Thompson CL, Pathak SD, Jeromin A, Ng LL, MacPherson CR, Mortrud MT, et al. Genomic anatomy of the hippocampus. Neuron. 2008;60(6):1010–21. doi: 10.1016/j.neuron.2008.12.008. PubMed PMID: 19109908.

23. Lu L, Igarashi KM, Witter MP, Moser EI, Moser MB. Topography of Place Maps along the CA3-to-CA2 Axis of the Hippocampus. Neuron. 2015;87(5):1078–92. doi: 10.1016/j.neuron.2015.07.007. PubMed PMID: 26298277.

24. Knierim JJ, Neunuebel JP. Tracking the flow of hippocampal computation: Pattern separation, pattern completion, and attractor dynamics. Neurobiology of learning and memory. 2016;129:38–49. doi: 10.1016/j.nlm.2015.10.008. PubMed PMID: 26514299; PubMed Central PMCID: PMC4792674.

25. Gore BB, Soden ME, Zweifel LS. Manipulating gene expression in projection-specific neuronal populations using combinatorial viral approaches. Current protocols in neuroscience. 2013;65:4 35 1–20. doi: 10.1002/0471142301.ns0435s65. PubMed PMID: 25429312; PubMed Central PMCID: PMC4242517.

26. Hnasko TS, Perez FA, Scouras AD, Stoll EA, Gale SD, Luquet S, et al. Cre recombinase-mediated restoration of nigrostriatal dopamine in dopamine-deficient mice reverses hypophagia and bradykinesia. Proceedings of the National Academy of Sciences of the United States of America. 2006;103(23):8858–63. doi: 10.1073/pnas.0603081103. PubMed PMID: 16723393; PubMed Central PMCID: PMC1466546.

27. Schwarz LA, Miyamichi K, Gao XJ, Beier KT, Weissbourd B, DeLoach KE, et al. Viral-genetic tracing of the input-output organization of a central noradrenaline circuit. Nature. 2015;524(7563):88–92. doi: 10.1038/nature14600. PubMed PMID: 26131933; PubMed Central PMCID: PMC4587569.

28. Franklin KBJ, Paxinos G. Paxinos and Franklin’s The mouse brain in stereotaxic coordinates. Fourth edition. ed. 1 volume (unpaged) p.

29. Bienkowski MS, Bowman I, Song MY, Gou L, Ard T, Cotter K, et al. Integration of gene expression and brain-wide connectivity reveals the multiscale organization of mouse hippocampal networks. Nature neuroscience. 2018;21(11):1628–43. doi: 10.1038/s41593-018-0241-y. PubMed PMID: 30297807; PubMed Central PMCID: PMC6398347.

30. Cembrowski MS, Wang L, Lemire AL, Copeland M, DiLisio SF, Clements J, et al. The subiculum is a patchwork of discrete subregions. eLife. 2018;7. doi: 10.7554/eLife.37701. PubMed PMID: 30375971; PubMed Central PMCID: PMC6226292.

31. Ding SL, Yao Z, Hirokawa KE, Nguyen TN, Graybuck LT, Fong O, et al. Distinct Transcriptomic Cell Types and Neural Circuits of the Subiculum and Prosubiculum along the Dorsal-Ventral Axis. Cell reports. 2020;31(7):107648. doi: 10.1016/j.celrep.2020.107648. PubMed PMID: 32433957.

32. Soudais C, Laplace-Builhe C, Kissa K, Kremer EJ. Preferential transduction of neurons by canine adenovirus vectors and their efficient retrograde transport in vivo. FASEB journal: official publication of the Federation of American Societies for Experimental Biology. 2001;15(12):2283–5. doi: 10.1096/fj.01-0321fje. PubMed PMID: 11511531.

33. Senn V, Wolff SB, Herry C, Grenier F, Ehrlich I, Grundemann J, et al. Long-range connectivity defines behavioral specificity of amygdala neurons. Neuron. 2014;81(2):428–37. doi: 10.1016/j.neuron.2013.11.006. PubMed PMID: 24462103.

34. Li SJ, Vaughan A, Sturgill JF, Kepecs A. A Viral Receptor Complementation Strategy to Overcome CAV-2 Tropism for Efficient Retrograde Targeting of Neurons. Neuron. 2018;98(5):905–17 e5. doi: 10.1016/j.neuron.2018.05.028. PubMed PMID: 29879392.

35. Wall NR, Wickersham IR, Cetin A, De La Parra M, Callaway EM. Monosynaptic circuit tracing in vivo through Cre-dependent targeting and complementation of modified rabies virus. Proceedings of the National Academy of Sciences of the United States of America. 2010;107(50):21848–53. doi: 10.1073/pnas.1011756107. PubMed PMID: 21115815; PubMed Central PMCID: PMC3003023.

36. Wickersham IR, Lyon DC, Barnard RJ, Mori T, Finke S, Conzelmann KK, et al. Monosynaptic restriction of transsynaptic tracing from single, genetically targeted neurons. Neuron. 2007;53(5):639–47. doi: 10.1016/j.neuron.2007.01.033. PubMed PMID: 17329205; PubMed Central PMCID: PMC2629495.

37. Tsien JZ, Chen DF, Gerber D, Tom C, Mercer EH, Anderson DJ, et al. Subregion-and cell type-restricted gene knockout in mouse brain. Cell. 1996;87(7):1317–26. doi: 10.1016/s0092-8674(00)81826-7. PubMed PMID: 8980237.

38. Moser MB, Moser EI. Distributed encoding and retrieval of spatial memory in the hippocampus. The Journal of neuroscience: the official journal of the Society for Neuroscience. 1998;18(18):7535–42. PubMed PMID: 9736671; PubMed Central PMCID: PMC6793256.

39. Moser MB, Moser EI. Functional differentiation in the hippocampus. Hippocampus. 1998;8(6):608–19. doi: 10.1002/(SICI)1098-1063(1998)8:6<608::AID-HIPO3>3.0.CO;2-7. PubMed PMID: 9882018.

40. Vann SD, Brown MW, Erichsen JT, Aggleton JP. Fos imaging reveals differential patterns of hippocampal and parahippocampal subfield activation in rats in response to different spatial memory tests. The Journal of neuroscience: the official journal of the Society for Neuroscience. 2000;20(7):2711–8. PubMed PMID: 10729352; PubMed Central PMCID: PMC6772240.

41. Jarrard LE. Selective hippocampal lesions: differential effects on performance by rats of a spatial task with preoperative versus postoperative training. Journal of comparative and physiological psychology. 1978;92(6):1119–27. doi: 10.1037/h0077516. PubMed PMID: 755058.

42. Olton DS, Walker JA, Gage FH. Hippocampal connections and spatial discrimination. Brain research. 1978;139(2):295–308. doi: 10.1016/0006-8993(78)90930-7. PubMed PMID: 624061.

43. Anagnostaras SG, Gale GD, Fanselow MS. The hippocampus and Pavlovian fear conditioning: reply to Bast et al. Hippocampus. 2002;12(4):561–5. doi: 10.1002/hipo.10071. PubMed PMID: 12201641.

44. Lein ES, Hawrylycz MJ, Ao N, Ayres M, Bensinger A, Bernard A, et al. Genome-wide atlas of gene expression in the adult mouse brain. Nature. 2007;445(7124):168–76. doi: 10.1038/nature05453. PubMed PMID: 17151600.

45. Alexander AS, Nitz DA. Retrosplenial cortex maps the conjunction of internal and external spaces. Nature neuroscience. 2015;18(8):1143–51. doi: 10.1038/nn.4058. PubMed PMID: 26147532.

46. Hafting T, Fyhn M, Molden S, Moser MB, Moser EI. Microstructure of a spatial map in the entorhinal cortex. Nature. 2005;436(7052):801–6. doi: 10.1038/nature03721. PubMed PMID: 15965463.

47. Taube JS, Muller RU, Ranck JB Jr., Head-direction cells recorded from the postsubiculum in freely moving rats. II. Effects of environmental manipulations. The Journal of neuroscience: the official journal of the Society for Neuroscience. 1990;10(2):436–47. PubMed PMID: 2303852; PubMed Central PMCID: PMC6570161.

48. Knierim JJ, Kudrimoti HS, McNaughton BL. Place cells, head direction cells, and the learning of landmark stability. The Journal of neuroscience: the official journal of the Society for Neuroscience. 1995;15(3 Pt 1):1648–59. PubMed PMID: 7891125; PubMed Central PMCID: PMC6578145.

49. Kjelstrup KG, Tuvnes FA, Steffenach HA, Murison R, Moser EI, Moser MB. Reduced fear expression after lesions of the ventral hippocampus. Proceedings of the National Academy of Sciences of the United States of America. 2002;99(16):10825–30. doi: 10.1073/pnas.152112399. PubMed PMID: 12149439; PubMed Central PMCID: PMC125057.

50. Lubenov EV, Siapas AG. Hippocampal theta oscillations are travelling waves. Nature. 2009;459(7246):534–9. doi: 10.1038/nature08010. PubMed PMID: 19489117.

51. Patel J, Fujisawa S, Berenyi A, Royer S, Buzsaki G. Traveling theta waves along the entire septotemporal axis of the hippocampus. Neuron. 2012;75(3):410–7. doi: 10.1016/j.neuron.2012.07.015. PubMed PMID: 22884325; PubMed Central PMCID: PMC3427387.

52. Kjelstrup KB, Solstad T, Brun VH, Hafting T, Leutgeb S, Witter MP, et al. Finite scale of spatial representation in the hippocampus. Science. 2008;321(5885):140–3. doi: 10.1126/science.1157086. PubMed PMID: 18599792.

53. Maurer AP, McNaughton BL. Network and intrinsic cellular mechanisms underlying theta phase precession of hippocampal neurons. Trends in neurosciences. 2007;30(7):325–33. doi: 10.1016/j.tins.2007.05.002. PubMed PMID: 17532482.

